# AI-Guided Discovery of Novel SARS-CoV-2 PLpro Inhibitors: Accelerating Antiviral Drug Development in the Fight Against COVID-19

**DOI:** 10.1101/2023.04.05.535700

**Authors:** Will Spagnoli, Wayne R Danter

**Affiliations:** CTO, Phronesis AI; acting CSO, Phronesis AI

## Abstract

The global pandemic caused by SARS-CoV-2 has highlighted the urgent need for effective antiviral drugs. The papain-like protease (PLpro) is a key viral enzyme involved in the replication and immune evasion of SARS-CoV-2, making it a promising target for antiviral drug development. In this study, we employed an artificial intelligence (AI)-driven drug discovery platform, LIME, to generate novel inhibitors of the SARS-CoV-2 PLpro. LIME is based on generative language models that can generate diverse, valid, and synthetically accessible compounds. The LIME software was used to identify potential inhibitors with strong binding affinity and specificity to the target protein. The top 13 hit compounds were tested in vitro, and the top 5 inhibitors with strong binding affinity and specificity were selected for further analysis. A top candidate molecule, CSEMRS-1376, exhibited similar binding energies and structural similarities to the known SARS-CoV-2 PLpro inhibitor, XR8-89. Computational analysis of the absorption, distribution, metabolism, excretion, and toxicity (ADMET) profiles of the hit compounds and XR8-89 showed that both CSEMRS-1376 and XR8-89 demonstrated favorable ADMET profiles. Overall, the LIME software successfully identified several novel molecules, including CSEMRS-1376, with strong potential as a therapeutic agent against SARS-CoV-2. The study highlights the potential of AI-driven drug discovery platforms, such as LIME, to accelerate the drug development process and pave the way for more efficient and effective therapeutics.

## Introduction

The global pandemic caused by the severe acute respiratory syndrome coronavirus 2 (SARS-CoV-2) has led to an unprecedented public health crisis, necessitating the rapid development of novel therapeutic strategies to combat the virus (1). Early efforts in this regard were the focus on the Spike protein as a therapeutic target. Another key viral enzyme involved in the replication and immune evasion of SARS-CoV-2 is the papain-like protease (PLpro) (2). Targeting PLpro with effective small molecule inhibitors offers a promising approach for the development of antiviral drugs, which could help in managing the disease and mitigating its impact on global health (3,12,13).

In recent years, artificial intelligence (AI) has emerged as a powerful tool in drug discovery, accelerating the process of designing and optimizing novel molecular structures with desirable pharmacological properties (4). AI-driven platforms have demonstrated their potential in a variety of applications, including the identification of potent inhibitors for various protein targets (5). One of the main advantages of using AI in drug discovery is the ability to rapidly generate and assess large libraries of molecules, making it possible to explore a vast chemical space in a cost-effective and time-efficient manner (6).

In this study, we leverage our proprietary Generative Language models (GLM) to generate molecular structures for synthesis and testing against the SARS-CoV-2 PLpro enzyme. Our aim is to design successful molecules that are safe, possess favorable absorption, distribution, metabolism, excretion, and toxicity (ADMET) properties, and effectively inhibit the PLpro enzyme, thus contributing to the development of potential therapeutic agents against SARS-CoV-2.

Our proprietary GLM are a type of machine learning model trained with large amounts of data to generate text by predicting the next word or character in a sequence. These models have demonstrated remarkable capabilities in natural language processing, text generation, and various other applications. In the context of drug discovery, GLM can be utilized to generate novel molecular structures which can be represented as SMILES strings. By training on large datasets of existing drug-like molecules, these models can produce diverse, valid, and synthetically accessible compounds. This approach enables the exploration of vast chemical spaces, potentially leading to the discovery of innovative therapeutics with desired properties, thereby revolutionizing the drug discovery process.

## Methods

The LIME Software and Drug Design Process: The LIME (Learning Iteratively through Molecular Evolution) software was employed to generate de novo inhibitors of the SARS-CoV-2 papain-like protease (PLpro) (Table 1). The input PDB file (7LBR) was based on the X-ray diffraction data from the study that developed the known PLpro inhibitor (XR8-89) (2). The LIME software was used to identify potential inhibitors with strong binding affinity and specificity to the target protein.

Table 1: The LIME de novo drug design process consists of the following steps:

1. An initial subset of structural starting points is chosen from Phronesis AI’s proprietary database.
2. These molecules are analyzed automatically using optimized virtual screening software.
3. The molecules and their detailed analysis are saved to the database and used for subsequent training.
4. The top-scoring molecules are selected based entirely on user-defined design objectives.
5. Novel molecules are produced by fine-tuning the molecular generation process.
6. The novel molecules are iteratively screened and added to the database alongside known molecules.
7. This optimization process is repeated autonomously and completed over the course of approximately 3-5 days.

As part of the design process, chemical synthetic feasibility, number of synthetic steps, and synthetic precursors are also predicted using AI-enabled retrosynthetic analysis to ensure the synthetic accessibility of all novel compounds generated by LIME. This approach has provided significant improvements over many alternative generative AI applications for drug design by preventing the LIME AI model from “cheating” and breaking chemical constraints to produce unrealistic molecules during its attempt to perform multi-attribute optimization of complex molecular structures.

LIME’s iterative design process yielded a focused molecular library of roughly ^~^10,000 novel drug candidates along with their respective virtual screening results for AutoDock Vina [14] predicted binding energy, AIZynthFinder AI-enabled retrosynthetic analysis [15], and cheminformatic molecular descriptors calculated using RDKit Open Source Cheminformatics Software [16]. As an additional virtual screening measure, we also performed in-silico analysis of all compounds within this focused drug library using Schrodinger Glide [17], our own implementation of the DeepDTA AI model [18] for protein-ligand affinity prediction screening, as well as our own proprietary affinity prediction AI model.

To further reduce the time and costs of drug synthesis, we then performed a substructure similarity search of our top-scoring drug candidates through a large compound library of molecules within the chemical spaces in which our synthetic chemists at OTAVA LTD specialize in. This allowed us to reduce the time and cost required for both acquiring the necessary synthetic precursors and performing the synthesis, while maintaining the optimized substructures and molecular properties identified by LIME’s design process. The top ten molecules were selected from LIME’s focused compound library based on a factor of their in silico predicted binding scores across the various screening software mentioned above, and these ten compounds were used for the substructure similarity search. All molecules within OTAVA LTD’s (https://otavachemicals.com) compound library having a substructure similarity greater than or equal to 0.7 with one of more of the ten selected molecules were aggregated to form a new cost-effective, focused compound library. This new focused library then was screened in-silico using the same methods, and the top 54 compounds were selected for in-vitro analysis based on factors of predicted binding, ADMET properties, and synthetic costs.

In-vitro testing: The 54 candidates identified by LIME underwent in-vitro testing using the BLI (Biolayer Interferometry) assay (OTAVA LTD) to assess their binding affinity to the SARS-CoV-2 PLpro. Of the top 13 candidates, the top 5 inhibitors with the best binding affinity and specificity were selected for further analysis. The basic elements of the BLI assay are presented in Figure 1.

**Figure 1:**
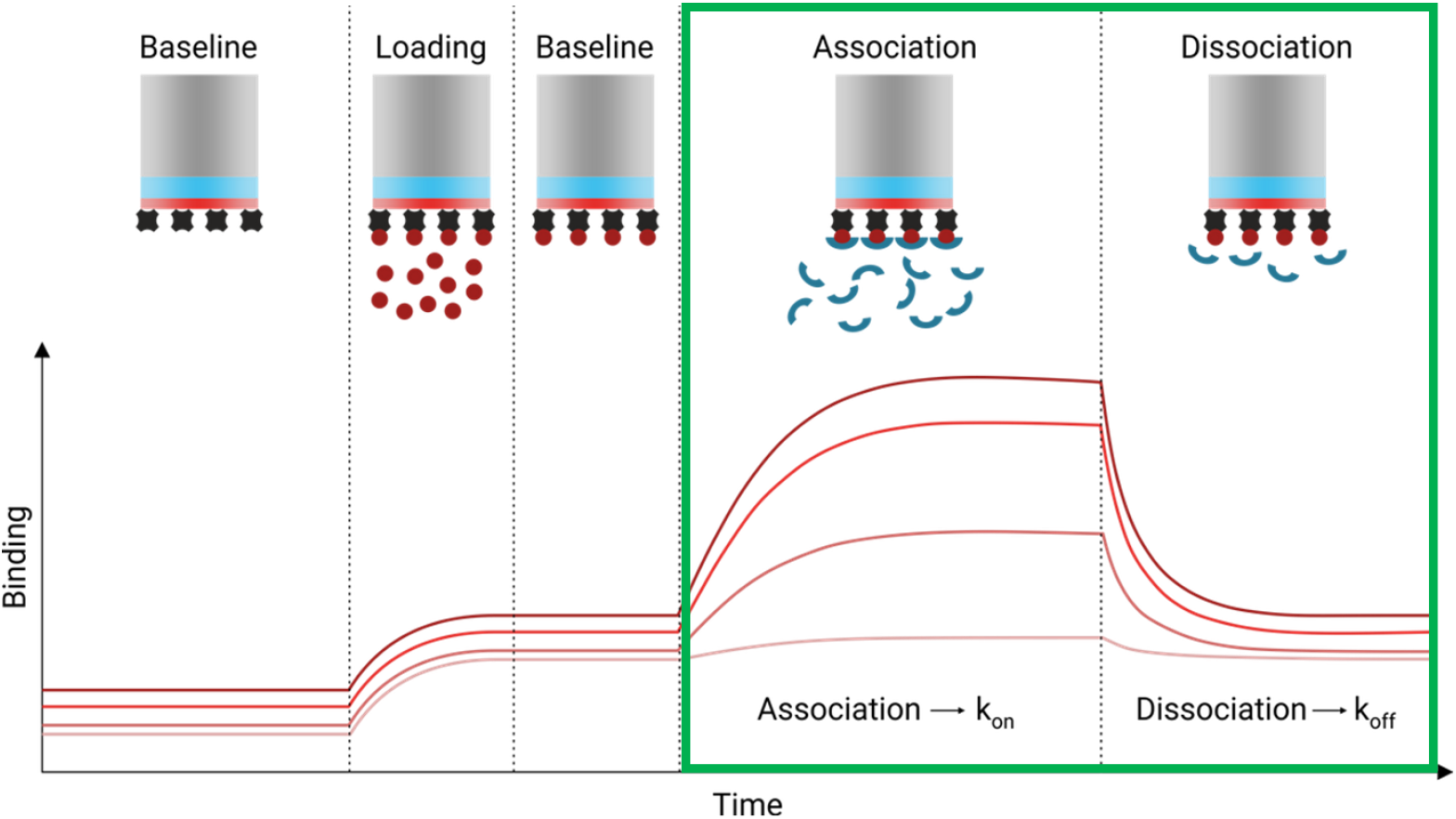
Overview of the BLI Binding interferometry assay used by OTAVA. The important components of the BLI interferometry binding assay. For the purposes of this study the important components are the Association phase from baseline followed by the Dissociation phase (outlined in **GREEN**).

### Molecular Docking and Binding Energy Analysis

AutoDock Vina was used to evaluate 9 different poses for one of the top candidate molecules, CSEMRS-1376, and the control molecule, XR8-89. Binding energy data arrays were obtained, and statistical analysis was conducted to compare the binding energies and determine the correlation between the two molecules.

### Visual Analysis

The binding modes and structural similarities of the top 5 hit compounds to the control molecule (XR8-89) were visually analyzed and compared to evaluate the performance of LIME in generating novel inhibitors.

### ADMET Profile Analysis

Computational analysis of the Absorption, Distribution, Metabolism, Excretion, and Toxicity (ADMET) profiles of the top 13 hit compounds and the known PLpro inhibitor (XR8-89) was performed using SWISSADME software (7) and DataWarrior/OSIRIS (8).

## Results

### The In vitro BLI Binding Study

Of the 54 candidates evaluated in vitro, 21 demonstrated potentially good/acceptable binding affinity but relatively poor specificity. Thirteen of these 21 candidates had modest/acceptable binding and variable selectivity and 5 modest/good binders had promising specificity. The binding results are presented in Figure 2 below.

**Figure 2:**
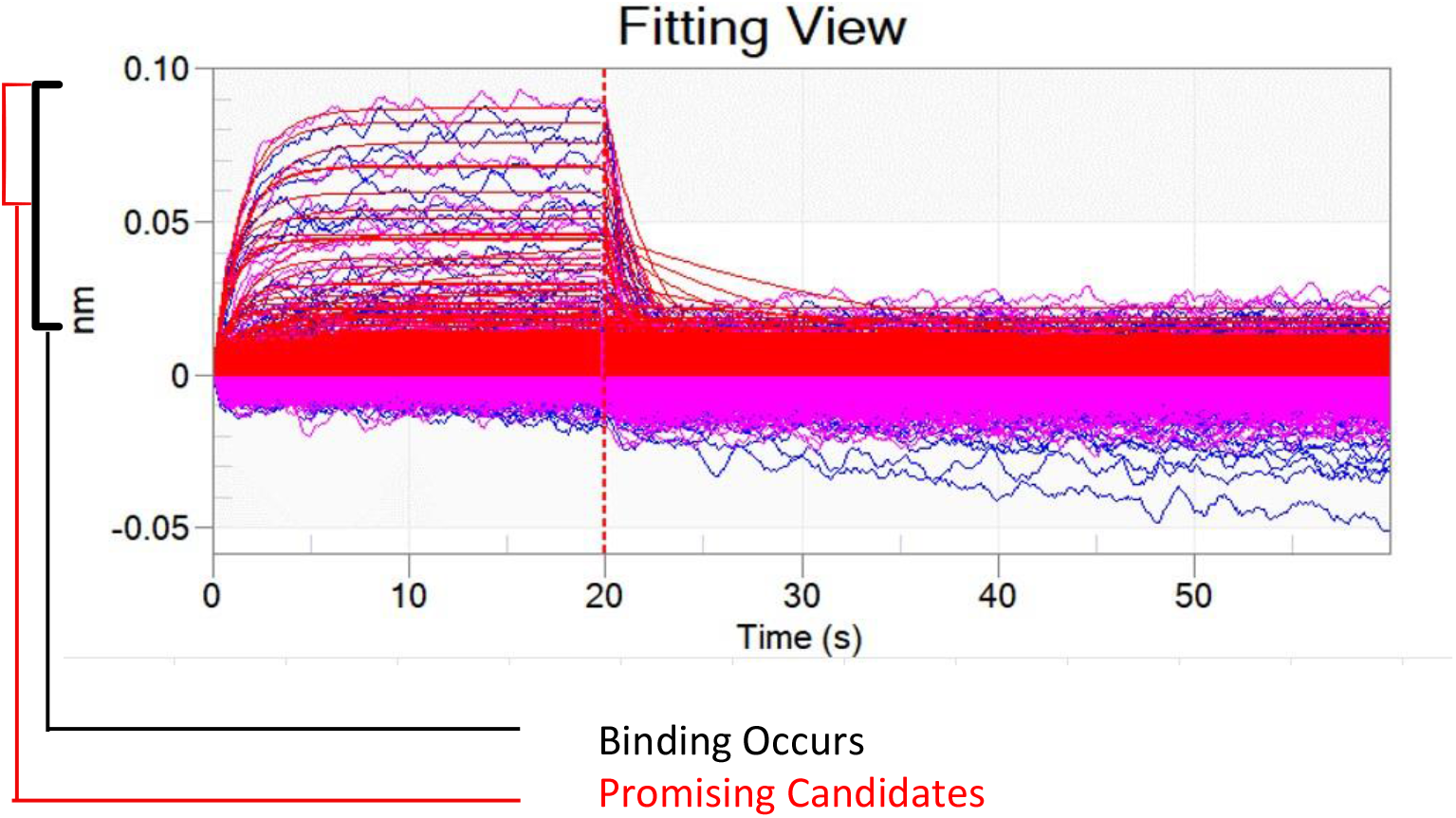
Raw Binding Data from BLI In Vitro assay for 54 test compounds. The BLI (Biolayer Interferometry*) assay measures direct binding between the SARS-CoV-2, PLpro enzyme and the 54 test compounds identified by the LIME AI. *Light is passed through samples. If binding occurs the wavelength of the light is distorted.

Comparative Binding Energy Analysis: The Pearson correlation coefficient between the binding energy data arrays of CSEMRS-1376 and XR8-89 was found to be 0.901, indicating a strong positive correlation. The unpaired t-test and Mann-Whitney U test showed no significant difference between the two groups.

#### Visual Analysis

A top candidate molecule, CSMERS-1376 was selected from the HITs as the focus of the blind test analysis because it exhibited the highest chemical and binding mode similarity to the control molecule (XR8-89) and it was also commercially available.

#### ADMET Profile Analysis

Both CSEMRS-1376 and XR8-89 demonstrated favorable ADMET profiles based on the computational analysis. While CSEMRS-1376 displayed a slightly lower molecular weight and higher topological polar surface area than XR8-89, its solubility was estimated to be lower. Both molecules were predicted to have high gastrointestinal absorption and good bioavailability, with CSEMRS-1376 not being a P-glycoprotein (Pgp) substrate, unlike XR8-89.

Overall, the LIME software successfully identified novel molecules, like CSEMRS-1376, with similar binding energies and structural similarities to the known SARS-CoV-2 PLpro inhibitor, XR8-89. Further in-vitro evaluation of the top 5 HITs, including CSEMRS-1376, is warranted to explore their potential as therapeutic agents against SARS-CoV-2.

## Discussion

In this study, we utilized the LIME software, an AI platform based on GLM, to generate de novo inhibitors of the SARS-CoV-2 papain-like protease (PLpro). The success of the LIME software in identifying novel molecules, including CSEMRS-1376, with strong binding affinity and specificity to the PLpro protein demonstrates the potential of GLM in the field of AI-driven drug discovery.

The rapid and efficient identification of novel drug candidates targeting specific binding pockets is crucial for accelerating the drug development process. GLM, like those employed by the LIME software, allow for the extraction of complex patterns and chemical features from large datasets, enabling the software to generate molecules with high structural similarity to known inhibitors (2). In the case of CSEMRS-1376, the LIME software was able to identify crucial substructures involved in protein-ligand binding while exploring variations in non-crucial substructures, thus providing a balance between maintaining key binding properties and exploring novel chemical space.

The use of AI-driven drug discovery platforms, such as LIME, offers several advantages over traditional drug discovery methods. These advantages include the ability to rapidly generate and screen large numbers of novel drug candidates, reducing the time and resources required for hit identification and lead optimization (3). Additionally, AI platforms can incorporate various aspects of drug development, such as ADMET profiling, into the molecule generation process, allowing for the identification of drug candidates with more favorable pharmacokinetic and pharmacodynamic properties (7, 8).

Despite the promising results obtained in this study, it is important to recognize the current limitations of AI-driven drug discovery platforms. One limitation is the reliance on accurate and comprehensive input data to ensure the generation of meaningful predictions. In the case of LIME, the success of the software in identifying CSEMRS-1376 and other HITs as potential PLpro inhibitors is contingent on the quality of the input PDB file and the annotation of the active site. Furthermore, while AI platforms can provide valuable insights into the potential binding affinity and ADMET properties of novel drug candidates, in vitro and in vivo validation is still necessary to confirm the efficacy and safety of these molecules.

In conclusion, this study highlights the potential of AI platforms, such as LIME, in the context of AI-driven drug discovery. The successful identification of CSEMRS-1376 and other HITs as potential PLpro inhibitors demonstrates the power of GLM in generating novel drug candidates with favorable binding properties and ADMET profiles. As the field of AI drug discovery continues to evolve, it is likely that the integration of AI platforms like LIME into the drug development process will become increasingly common, paving the way for more rapid, efficient, and effective targeted therapeutics.

